# Examining Fetal Sex-Specific Placental DNA Methylation Intensities and Variability Post *In Vitro* Fertilization

**DOI:** 10.1101/2024.08.08.604307

**Authors:** Melanie Lemaire, Keaton Warrick Smith, Samantha L Wilson

## Abstract

Infertility impacts up to 17.5% of reproductive-aged couples worldwide. To aid in conception, many couples turn to assisted reproductive technology, such as *in vitro* fertilization (IVF). IVF can introduce both physical and environmental stressors that may alter DNA methylation regulation, an important and dynamic process during early fetal development. This meta-analysis aims to assess the differences in the placental DNA methylome between spontaneous and IVF pregnancies. We identified three studies from NCBI GEO that measured DNA methylation with an Illumina Infinium Microarray in post-delivery placental tissue from both IVF and spontaneous pregnancies with a total of 575 samples for analysis (n = 96 IVF, n = 479 spontaneous). While there were no significant or differentially methylated CpGs in mixed or female stratified populations, we identified 9 CpGs that reached statistical significance (FDR <0.05) between IVF (n = 56) and spontaneous (n = 238) placentae. 7 autosomal CpGs and 1 X chromosome CpG was hypermethylated and 2 autosomal CpGs were hypomethylated in the IVF placentae compared to spontaneous. Autosomal CpGs closest to *LIPJ*, *EEF1A2*, and *FBRSL1* also met our criteria to be classified as biologically differentially methylated CpGs (FDR <0.05, |Δ*β*|>0.05). When analyzing variability differences in Δ*β* values between IVF females, IVF males, spontaneous females and spontaneous males, we found a significant shift to greater variability in the both IVF males and females compared to spontaneous (p <2.2e-16, p <2.2e-16). Trends of variability were further analyzed in the biologically differentially methylated autosomal CpGs near *LIPJ EEF1A2*, and *FBRSL1*, and while these regions were statistically significant in males, the female Δ*β*s and ΔCoVs followed a similar trend that differed in magnitude. In males and females there was a statistically significant difference in proportions of endothelial cells, hofbauer cells, stromal cells and syncytiotrophoblasts between spontaneous and *in vitro* Fertilization (IVF) populations. We also observed significant differences between sex within reproduction type in syncytiotrophoblasts and trophoblasts. The results of this study are critical to further understand the impact of IVF on tissue epigenetics which may help to investigate the connections between IVF and negative pregnancy outcomes. Additionally, our study supports sex specific differences in placental DNA methylation and cell composition should be considered as factors for future placental DNA methylation analyses.

## Introduction

In the last 30 years, infertility rates in reproductive-aged couples have increased from 8-12% up to roughly 17.5%^1,2^. Rising infertility rates are partly linked to the trend of later motherhood in many developed countries, driven by higher levels of maternal education, increased female employment, and financial uncertainty^3,4^. Assisted reproductive technology (ART) encompasses many medical resources that aid in conception for those struggling with infertility, including those of delayed reproductive age, couples in same-sex relationships, those wishing to use monogenic screening, and those opting for frozen or donor oocytes. With a growing demand for ART, investigation of the impact of ART techniques on maternal and fetal health is crucial to ensure a standard level of care for all those in need of reproductive assistance.

ART procedures can include *in vivo* techniques such as intrauterine insemination, or *in vitro* techniques such as IVF or Intracytoplasmic Sperm Injection (ICSI)^5^. Currently, IVF is the most commonly used ART^6,7^, with over 8 million children born from IVF in the last 40 years^8^. In the process of IVF, oocytes are retrieved, typically following a period of controlled ovarian stimulation with the use of exogenous gonadotropins, and are fertilized by donor sperm in media that mimics the *in vivo* reproductive environment^6,9^. Embryos are incubated until the cleavage stage (day 3) or blastocyst stage (day 5), with the latter more common, before transfer to the uterus^6^.

Despite IVF’s benefits in aiding conception, its procedure introduces additional stressors into early development that are not observed in spontaneously conceived pregnancies. Ovarian stimulation and subsequent oocyte retrieval contribute both hormonal and physical disturbances^10,11^. Embryo incubation conditions including oxygen concentration, pH, temperature, and osmolarity are highly dynamic within the reproductive tract and difficult to replicate in an *in vitro* setting. Estimated measurements of these biological conditions are mainly from animal studies, which may not be optimal for human development^9,12^. Other factors such as fresh or frozen embryo transfer and use of preimplantation genetic testing (PGT) may further contribute to the diversity of stressors IVF may impose on early development.

Current literature has connected ART techniques, including IVF, with an increased risk of many negative pregnancy outcomes. Rates of preterm birth, major birth defects, preeclampsia, cardiovascular issues, and imprinting disorders such as Beckwith-Wiedemann syndrome, have all been reported at higher proportions in ART conceived pregnancies^13–16^. However, inter-study results in the literature are contradictory, and it is difficult to parse through if these effects are due to the ART protocols or underlying infertility, as a high proportion of those seeking ART are doing so for female and or male factor infertility^7^.

It has been further hypothesized that epigenetic factors may play a role in this observed relationship between IVF and an increased risk of negative pregnancy outcomes. Epigenetic modifications are heritable and may alter gene expression without changing the DNA sequence, through processes such as histone modifications, non-coding RNA interference, and DNA methylation^17,18^.

DNA methylation occurs through the addition of a methyl group to the fifth carbon of a cytosine that precedes a guanine, typically referred to as a cytosine guanine dinucleotide (CpG)^18,19^. This mechanism is important for regulating transcription in processes such as X inactivation in XX individuals, cell differentiation, and genomic imprinting^20^. Both the establishment of DNA methylation and de-methylation can occur dynamically, providing a level of plasticity that may be influenced by environmental factors. However, the extent by which DNA methylation contributes to the interaction between the environment and gene regulation remains unknown^21^.

Dynamic epigenetic regulation is an important process in early embryo development. Following fertilization, there is a genome-wide loss of DNA methylation, aside from imprinted regions, prior to totipotency acquisition and cell differentiation^22,23^. Once the blastocyst stage is reached, re-methylation occurs with distinct and asynchronous patterns in the trophectoderm, which will differentiate into the main cell types of the placenta and primitive endoderm^22^. As the main steps of IVF occur during this window of DNA methylation reprogramming, it is questioned if exposure to *in vitro* conditions may negatively impact this process. Therefore the connection between IVF stressors and an increased risk of negative pregnancy outcomes may be due to IVF’s impact on DNA methylation in early development which may lead to gene dysregulation in cell types crucial for placental health.

Assessment of placental DNA methylation patterns following IVF pregnancies compared to spontaneous pregnancies may give insight to the effects of IVF on placental development. Recent meta-analyses have qualitatively compared the results of studies that analyzed DNA methylation data from ART pregnancies in various tissues^22,24^. One limitation of these meta-analyses is many of the included studies used fundamentally different techniques for normalization and statistical analyses in their investigation of DNA methylation. In our study we aim to combine and reevaluate raw publicly available DNA methylation datasets from placental tissue following IVF or spontaneous conception in a quantitative meta-analysis. We hypothesize that DNA methylation levels will be different between IVF and spontaneously conceived pregnancies.

## Methods

### Dataset Collection

This study adheres to the guidelines outlined in the Preferred Reporting Items for Systematic Reviews and Meta-Analyses (PRISMA) 2020 statement^25^. Potential datasets were identified by searching NCBI Gene Expression Omnibus (GEO) using keywords related to *in vitro* fertilization in human participant studies. Data sets were first selected on the basis of their titles and abstracts. Datasets included after first round screening underwent full paper review. Studies were included if they measured DNA methylation from both IVF and spontaneously conceived healthy, full-term, singleton pregnancies. Full-term placenta samples were selected to ensure tissue sampling and clinical data collection did not occur prior to development of pregnancy complications that may introduce pathology specific placental DNA methylation patterns. Preterm placentas were excluded as they are known to have different placental DNA methylation profiles compared to full-term controls^26^. Studies were excluded if they did not report both IVF and spontaneous samples, were from pregnancies with only multiples, or only had samples from complicated or premature pregnancies. A flowchart showing inclusion and exclusion screening decisions is shown in Figure 1. All screening steps were performed by one researcher (ML).

**Figure 1:**
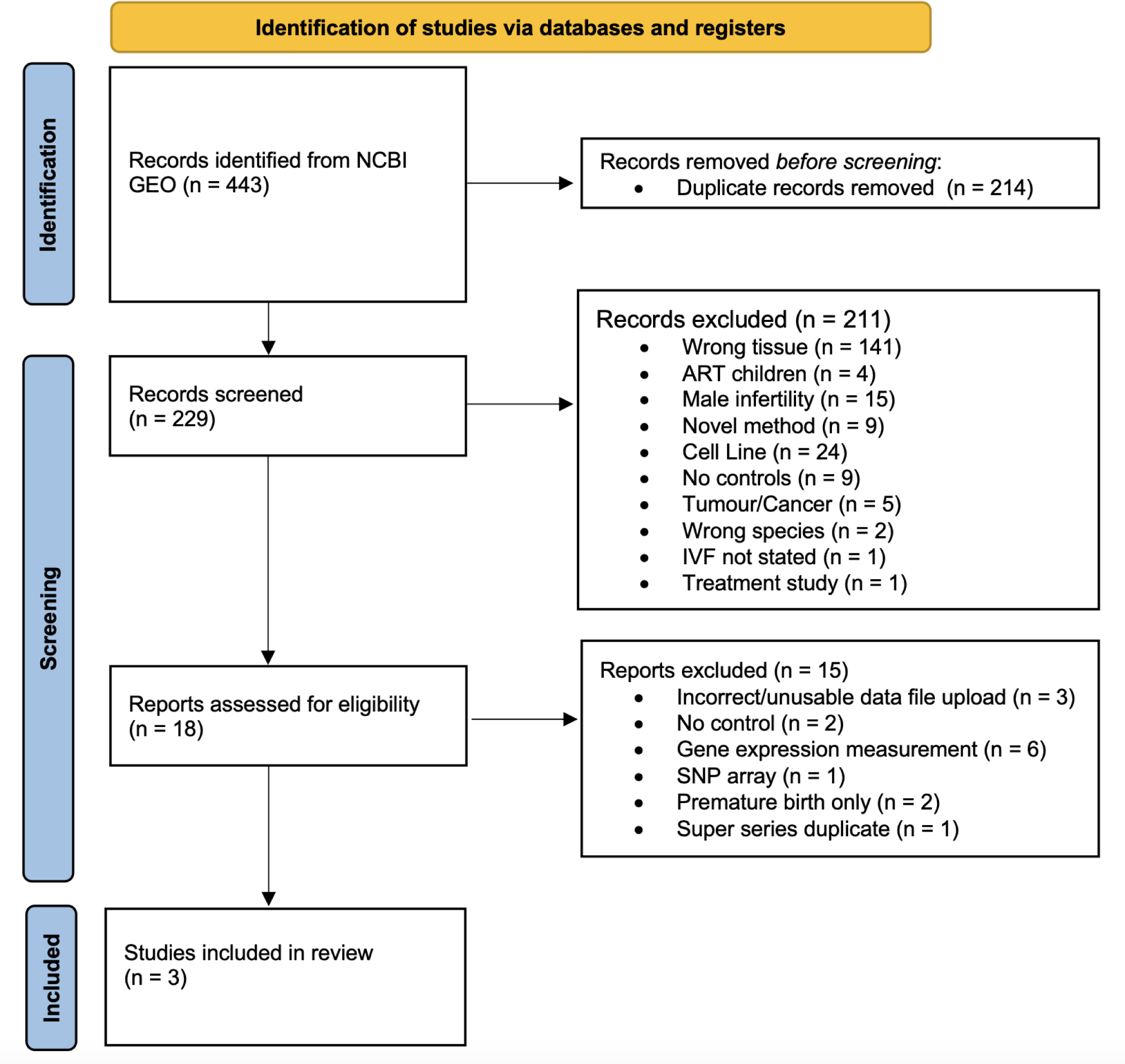
Flowchart of dataset selection based on inclusion and exclusion criteria, assembled using the PRISMA 2020 guidelines.

Screening NCBI GEO revealed 3 studies that had raw DNA methylation files measured by either Illumina Infinium 450K or EPIC bead chips in placental tissue collected at time of delivery from singleton, uncomplicated, full-term pregnancies (GSE120250^27^, GSE75248^28^, GSE208529^29^). Each study reported conception type, fetal sex, sentrix position, and sentrix ID and had raw intensity data (IDAT) files available for each sample.

### Data Quality Control and Normalization

R version 4.4.1^30^ was used for all subsequent data processing and analysis. All clinical and technical information reported for each study sample was extracted using the GEOquery package^31^ version 2.72.0. IDAT files containing intensities for each probeset on Illumina 450K and EPIC arrays were downloaded from NCBI GEO for each included dataset onto a server. As a quality control step to ensure each sample reflected the reported clinical project data, we compared reported fetal sex to fetal sex predictions. Fetal sex was predicted based on DNA methylation intensities from the XY chromosomes using the minfi package^32^ version 1.50.0. Samples that did not match the reported and predicted fetal sex were excluded (n = 5), as it could not be determined if these samples were mislabeled during the collection process. Samples within each dataset that used *in vivo* techniques (n = 21) such as intrauterine insemination, were maternal facing placental samples (n = 195), did not report conception type (n = 9), or were specified as outliers (n = 9) were removed from the datasets. This left a combined sample size of 575 for analysis (n = 96 IVF, n = 479 Spontaneous) as shown in Table 1. Background correction and normalization of DNA methylation intensities was implemented using adjustedFunnorm^33^ through the wateRmelon package version 2.11.2.

**Table 1:**
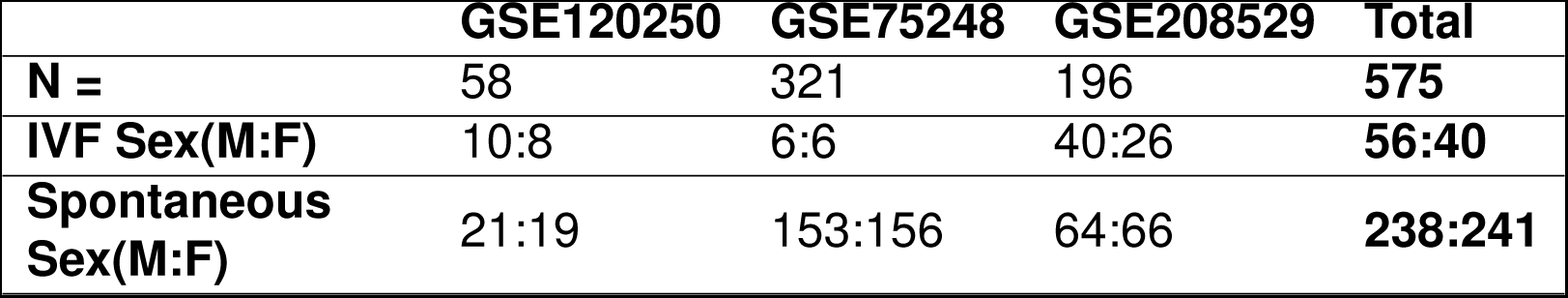
Sample demographics for each of the three GEO studies used in this meta-analysis.

### Probe Filtering

Probe filtering was completed on X and Y chromosomes and autosomes using a modified DNA methylation processing pipeline, to allow integration of X and Y chromosome data, as outlined by Inkster et.al. (2023). Bad quality probes defined as having a missing *β* value <0.05 or detection p-value <0.05 in >5% of samples were removed (n = 2189). Probes that bind to known Single Nucleotide Polymorphisms (SNP) regions (n = 19361) or that are known to crosshybridize to multiple sites (n = 36571) were removed as per the annotation from Price et.al. (2013). We filtered out non-variable placental CpG probes (n = 89160) based on the annotation from Edgar et.al. (2017). After probe filtering, we analyzed a total of 296545 autosomal CpG’s and 8779 X chromosome CpG’s. An additional 227 Y chromosome CpG’s were analyzed in male samples.

### Differential DNA Methylation Analysis

DNA methylation intensities measured at individual probe-specific CpG’s were compared between IVF and spontaneous placental populations using linear regression models implemented through the limma package version 3.60.4^37^. This was first conducted in the mixed fetal sex population using a model that corrected for conception type, GEO study number, and fetal sex.

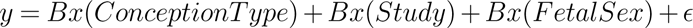

In fetal stratified populations we used a model that corrected for conception type and GEO study number only.

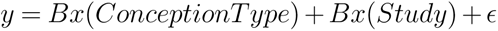

GEO study numbers, which connect samples to each of the 3 datasets, were used to mitigate batch effects that may be introduced when cross analyzing data from separate datasets together. We opted not to use ComBat, a common batch effect correction method, as previous studies reported that ComBat may increase false positives in unbalanced study designs and decrease reproducibility of findings^38,39^. As this meta-analysis compared studies that were not designed to balance each other, we wanted to limit the potential of ComBat introducing spurious findings, indicative of applying it to an unbalanced study design.

Following linear modelling, statistically significant CpGs were defined as having a False Discovery Rate (FDR) <0.05. Differentially methylated CpGs were defined as having a FDR <0.05 and a change in DNA Methylation (|Δ*β*|) >0.05. Δ*β*’s were calculated for each probe by subtracting the mean DNA methylation *β* value in the IVF population from the *β* value in the respective spontaneous population. Differentially methylated CpGs reaching the more stringent criteria are more likely to indicate a biologically meaningful result.

### Autosomal variability analysis

To analyze variability in autosomal DNA methylation intensities between populations, we calculated coefficients of variance (CoV) for measured *β* values across all samples in IVF males, IVF females, spontaneous males and spontaneous female populations. The densities of the CoV’s for each group were plotted and distributions were compared using a Kolmogorov-Smirnov (KS) test.

### Visualizing Key Probe Data on Genome Browser Tracks

Visualization of population CoV’s, Δ*β*’s between IVF and spontaneous in males and females, and ΔCoV’s between IVF and spontaneous in males and females were visualized and superimposed on a genome track for differentially methylated autosomal CpGs. Genomic coordinates of the methylation array probes were annotated using the HM450 hg19 manifest from the Children’s Hospital of Philadelphia Zhou Lab Infinium Annotation GitHub site^40^. For each group, coefficients of variance at the genomic coordinates of the probes were plotted using the Gviz version 1.48.0 library^41^ for R. Annotations for transcripts at or near the probes were obtained from the TxDb.Hsapiens.UCSC.hg19.knownGene package version 3.2.2^42^ and mapped to gene symbols using the EnsDb.Hsapiens.v75 package version 2.99.0^43^ for R. Coordinates for CpG islands were obtained from the UCSC GRCh37/hg19 assembly CpG island annotation table^44^.

### Placental Cell Deconvolution

Placental cell proportions were estimated using the planet package version 1.12.0 and EpiDISH package version 2.20.0 using placental reference CpGs^45^. Proportions of trophoblasts, stromal cells, hofbauer cells, endothelial cells, nucleated Red Blood Cells (nRBC), and syncytiotrophoblasts were calculated for each sample. An ANOVA with a Bonferonni test was performed to compare mean cell proportions between IVF Female, IVF male, spontaneous Female and spontaneous male populations for each individual cell type.

## Results

### Fetal male stratified population revealed differentially methylated autosomal CpGs between IVF and spontaneous placentae

We first assessed placental DNA methylation at individual autosomal CpG’s between IVF and spontaneously conceived pregnancies. Statistically significant CpGs were defined as having a FDR <0.05. Differentially methylated CpGs were defined as having a FDR <0.05 and a change in DNA Methylation (|Δ*β*|) >0.05 to denote potential biological significance.

In our full mixed-fetal sex population we identified no significant or differentially methylated CpGs between IVF (n = 96) and spontaneous (n = 479) placentae as shown in Figure 2A. Following full population analysis, we stratified out populations by fetal sex. Sex stratified analysis of DNA methylation in the fetal female population also revealed no significant or differentially methylated CpGs between IVF (n = 40) and spontaneous (n = 241) placentae as shown in Figure 2B. However, in the male stratified population we identified 9 CpGs that reached statistical significance between IVF (n = 56) and spontaneous (n = 238) placentae as shown in Figure 2C. 7 CpGs were hypermethylated in the IVF placentae compared to spontaneous with *LIPJ, C19orf25, MUM1, RASGRF1, FAF1, AK092451,* and *EEF1A2* as the closest transcriptional start sites. 2 CpGs was hypomethylated in the IVF placentae compared to spontaneous and with their closest transcriptional start sites of *EXD3* and *FBRSL1* as shown in Table 2. The CpGs closest to LIPJ, EEF1A2, and FBRSL1 also met our criteria to be classified as differentially methylated CpGs.

**Figure 2:**
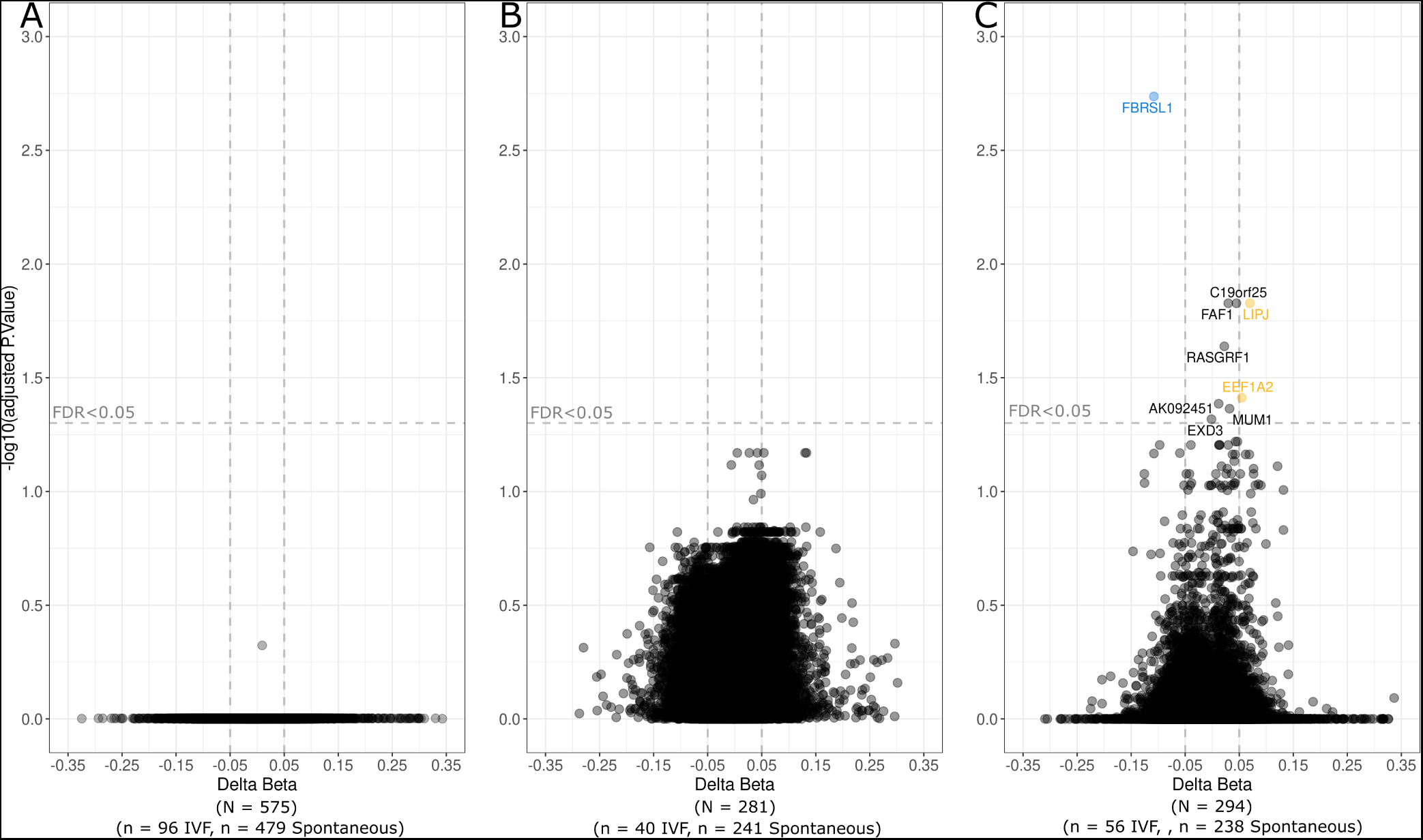
Volcano plots depicting differentially methylated autosomal sites between IVF and spontaneous placentae in (A) mixed fetal sex population, (B) female stratified population, and (C) male stratified population. –log10 of the adjusted p-value is plotted on the y axis and change in DNA methylation (Δ*β*) is plotted on the x axis. CpG’s highlighted in blue are differentially hypomethylated in IVF (Δ*β* <-0.5, FDR <0.05). CpGs highlighted in yellow are differentially hypermethylated in IVF (Δ*β* >0.5, FDR <0.05). Sample size is indicated under x-axis.

**Table 2:**
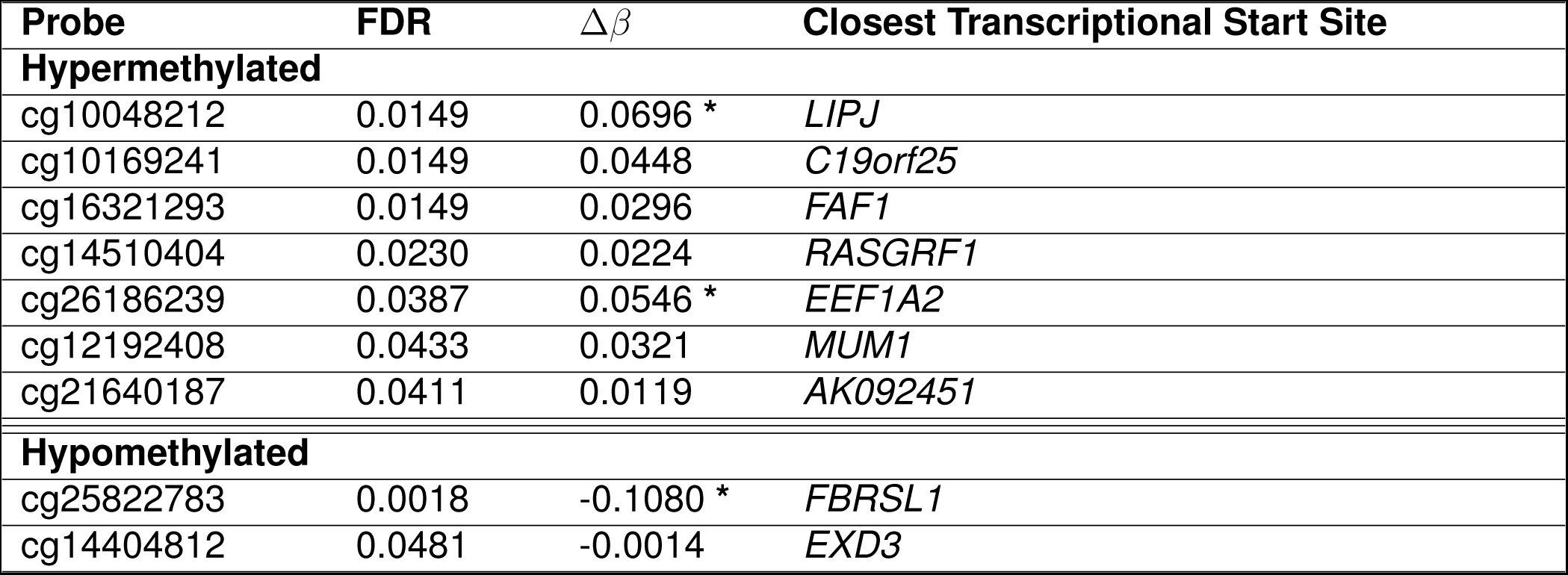
Autosomal probes with significantly different DNA methylation in IVF fetal male placentae compared to spontaneous identified from linear modelling analysis. Asterisks indicate Δ*β* values that met the differential methylation criteria to indicate potential biological significance

### No differentially methylated CpGs were identified in sex stratified analyses of XY chromosomes

We assessed DNA methylation in the X chromosome between IVF and spontaneous placentae in sex stratified populations. In the female population we observed no significant or differentially methylated CpGs between IVF and spontaneous placentae as shown in Figure 3A. In the male population we found one statistically significant CpG with a closest transcriptional start site to CAPN6 in Figure 3B. However this CpG did not reach our biological criteria to be considered differentially methylated as shown in Table 3

**Figure 3:**
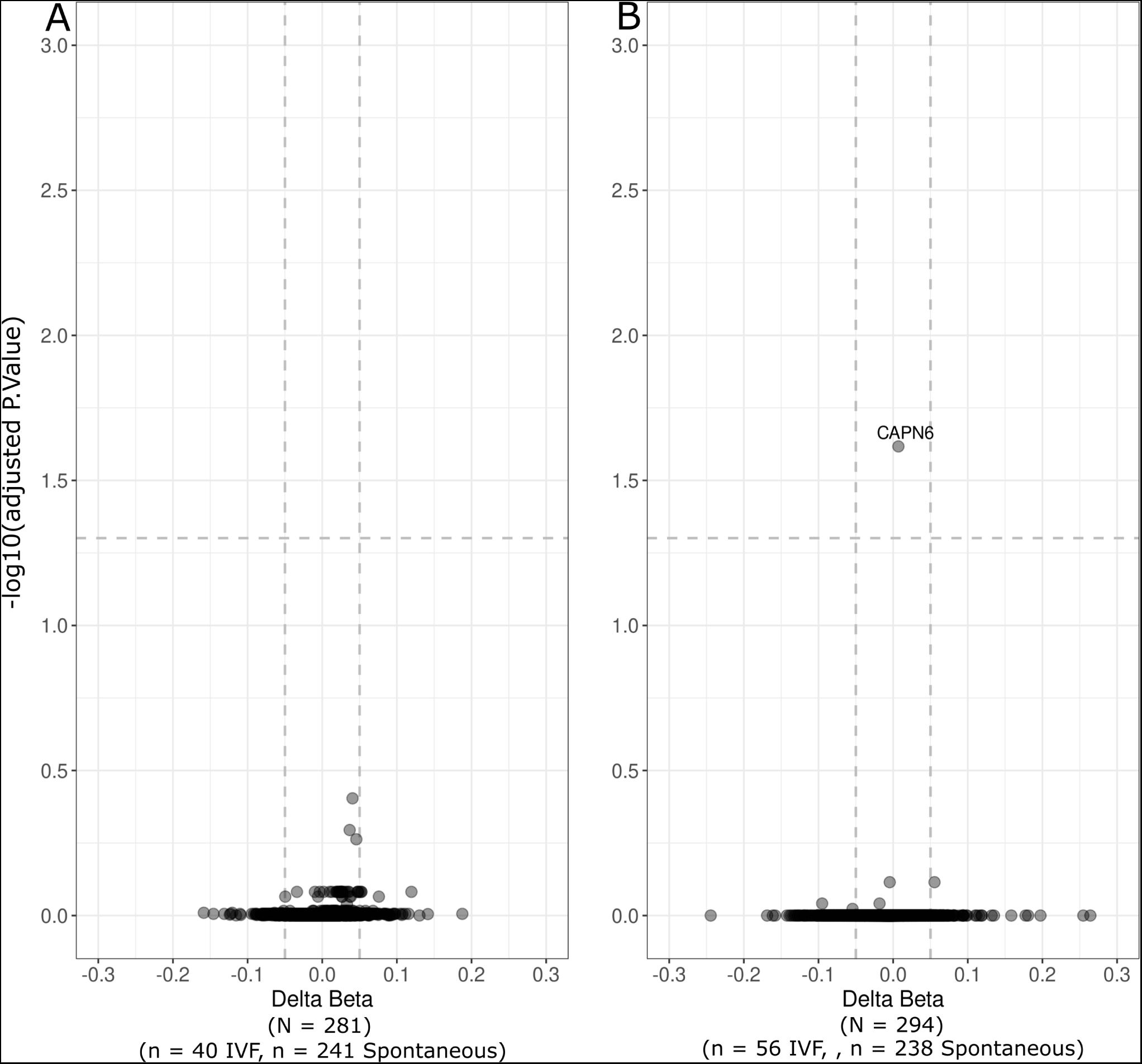
Volcano plots depicting X chromosome probe methylation differences between IVF and spontaneous placentae in (A) female stratified population and (B) male stratified population following linear modelling. –log10 of the adjusted p-value is plotted on the y axis and change in DNA methylation (Δ*β*) is plotted on the x axis.

**Table 3:**
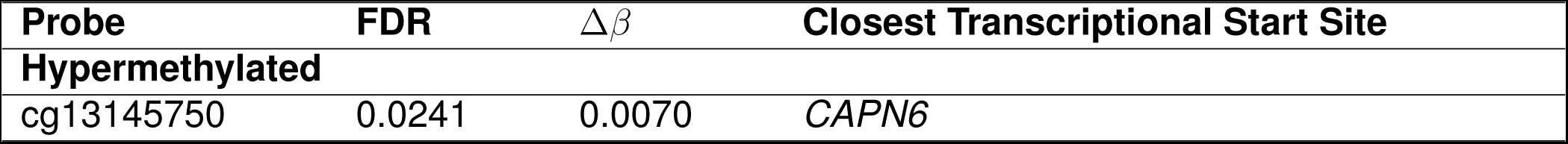
X chromosome probe with statistically different DNA methylation in IVF fetal male placentae identified from linear modelling analysis.

Probes with significantly different DNA methylation in IVF fetal male placentae compared to spontaneous identified from linear modelling analysis. Asterisks indicate Δ*β* values that met the differential methylation criteria to indicate potential biological significance

We assessed DNA methylation in the Y chromosome between IVF and spontaneous placentae in the male stratified population. No significant or differentially methylated CpGs were observed (Supplementary Figure 1).

### Variability in placental DNA methylation increased in IVF sex-stratified populations

Coefficients of variance were calculated for all samples in IVF male, IVF female, spontaneous male and spontaneous female populations and plotted in a density plot as shown in Figure 4 After comparing the densities of coefficients of variance between IVF male, IVF female, spontaneous male and spontaneous female populations we found that both male and female IVF populations had a significant shift in density towards larger CoV’s compared to spontaneous (P<2.2e-16, P<2.2e-16). IVF females show a slightly larger shift to higher CoV’s comapred to IVF males (P<2.2e-16).

**Figure 4:**
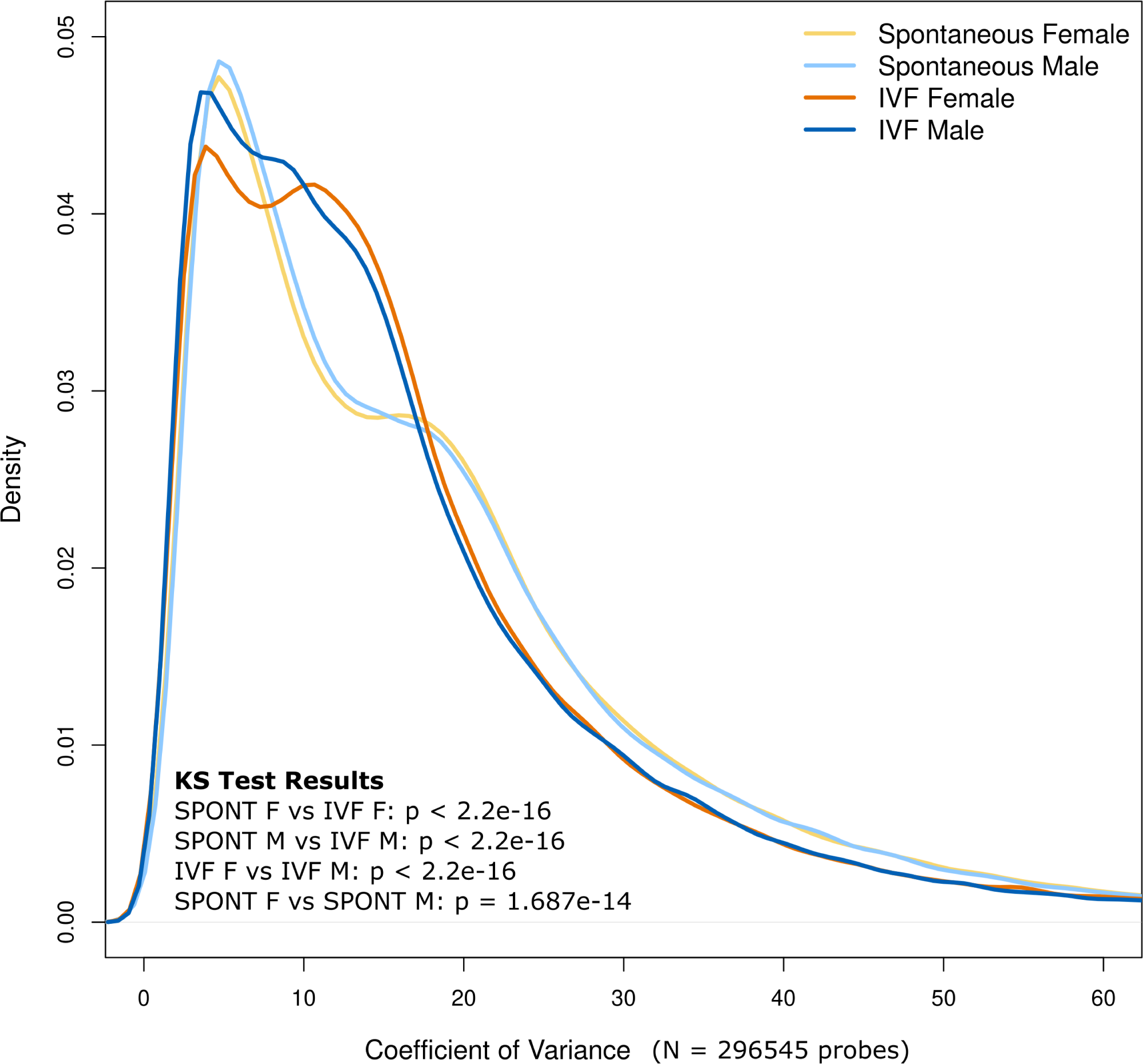
Distribution of Coefficients of Variance in sex stratified IVF and Spontaneous populations. Results from KS tests are denoted in the bottom left corner

CoV data from each population and male and female Δ*β*’s and ΔCoV’s comparing IVF to spontaneous were superimposed on genome tracks for each of the 3 differentially methylated autosomal CpGs identified in male IVF Placentae as shown in Figure 5. We observed consistent higher ΔCoV’s between IVF and spontaneous in male placentae in the regions of *LIPJ* and *FBRSL1* while the region of *EEF1A2* observed less of a male dominated increase in ΔCoV.

**Figure 5:**
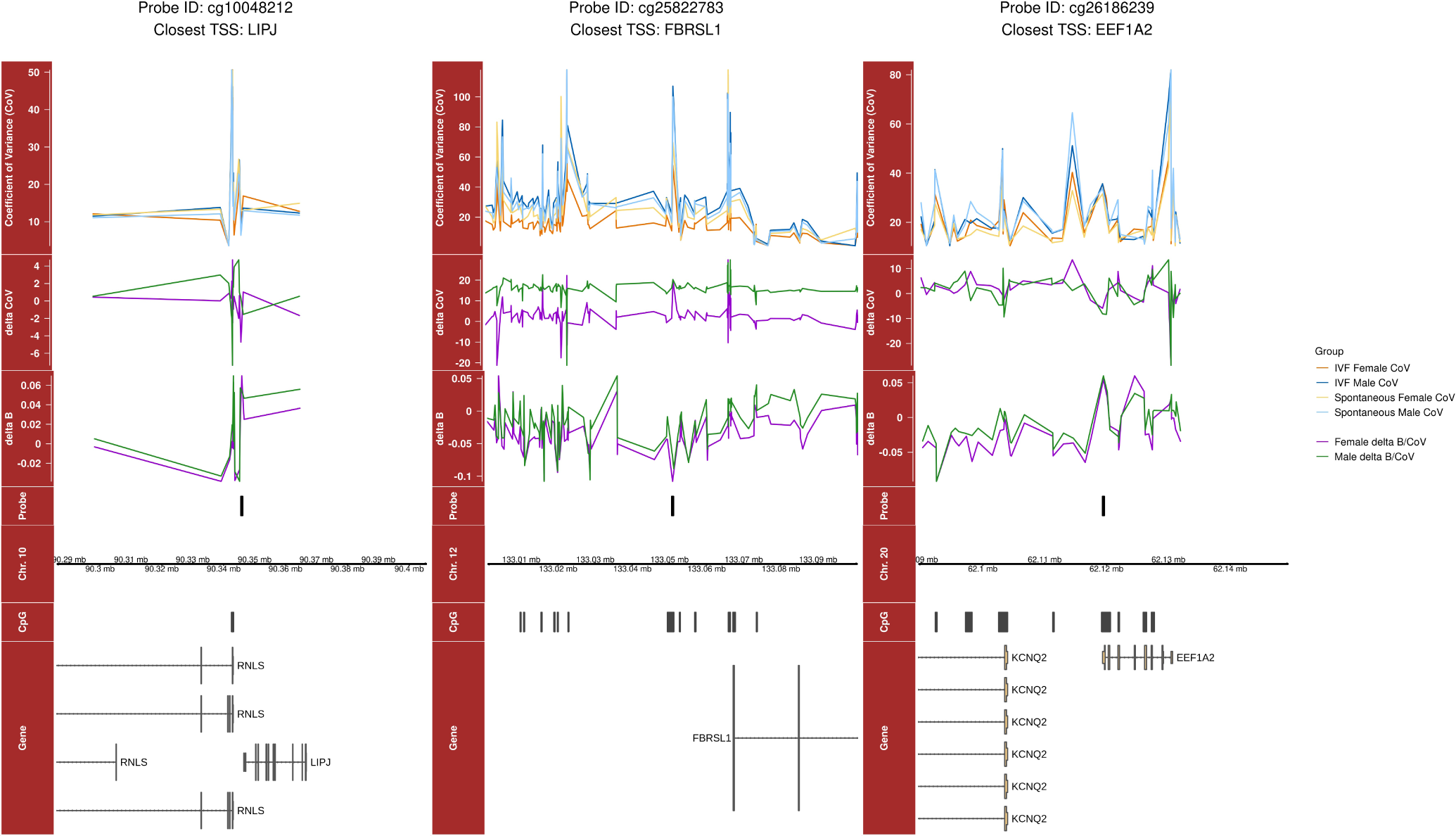
Coefficients of variance (CoV) for each of the four population groups as well as Δ*β* and ΔCoV for male and female samples are plotted for the genomic regions surrounding cg10048212, cg25822783, and cg26186239 methylation probes. Genomic coordinates of each probe are shown along with their position relative to CpG islands and nearby known genes annotated from their respected UCSC track and the Ensembl database. Plots were produced using the Gviz package for R.

Δ*β*’s followed a similar trend with higher Δ*β*’s between IVF and spontaneous in male placentae in the regions of *LIPJ* and *FBRSL1* while the region of *EEF1A2* was more comparable to Δ*β*’s of females. However, we noted that the overall patterns observed in ΔCoV’s and Δ*β*’s across each gene were similar between males and females, despite differences in the magnitudes of measured values. Genome tracks for the 6 significant but not differentially methylated autosomal CpGs identified in male IVF placentae are available in Supplementary Plots 2-7.

### Placental cell-type proportions differed between sex and conception type

We estimated placental cell-type composition in each of our samples using a placental cell deconvolution meathod. Using an ANOVA with a Bonferroni test we found that there were significantly different proportions of different cell-types within sex between IVF and spontaneous placenta, as well as within treatment between sex as shown in Figure 6. In females, proportions of endothelial cells, hofbauer cells, stromal cells, and syncytiotrophoblasts differed significantly between IVF and spontaneous populations (p = 5.1e-5, p = 0.0020, p = 1.5e-6, p = 0.0045 respectively). In males, proportions of endothelial cells, hofbauer cells, stromal cells, and synsytiotrophoblasts also differed significantly between IVF and spontaneous populations (p = 4.5e-9, p = 0.0028, p = 5.5e-6, p = 5.5e-6 respectively). We also found differences between sex within conception type. Males had a significantly higher proportion of synsytiotrophoblasts compared to females in both the IVF and spontaneous groups (p = 0.0419 and p = 0.021 respectively). Spontaneous males also had a significantly lower proportion of trophoblasts compared to spontaneous females (p = 0.0021)

**Figure 6:**
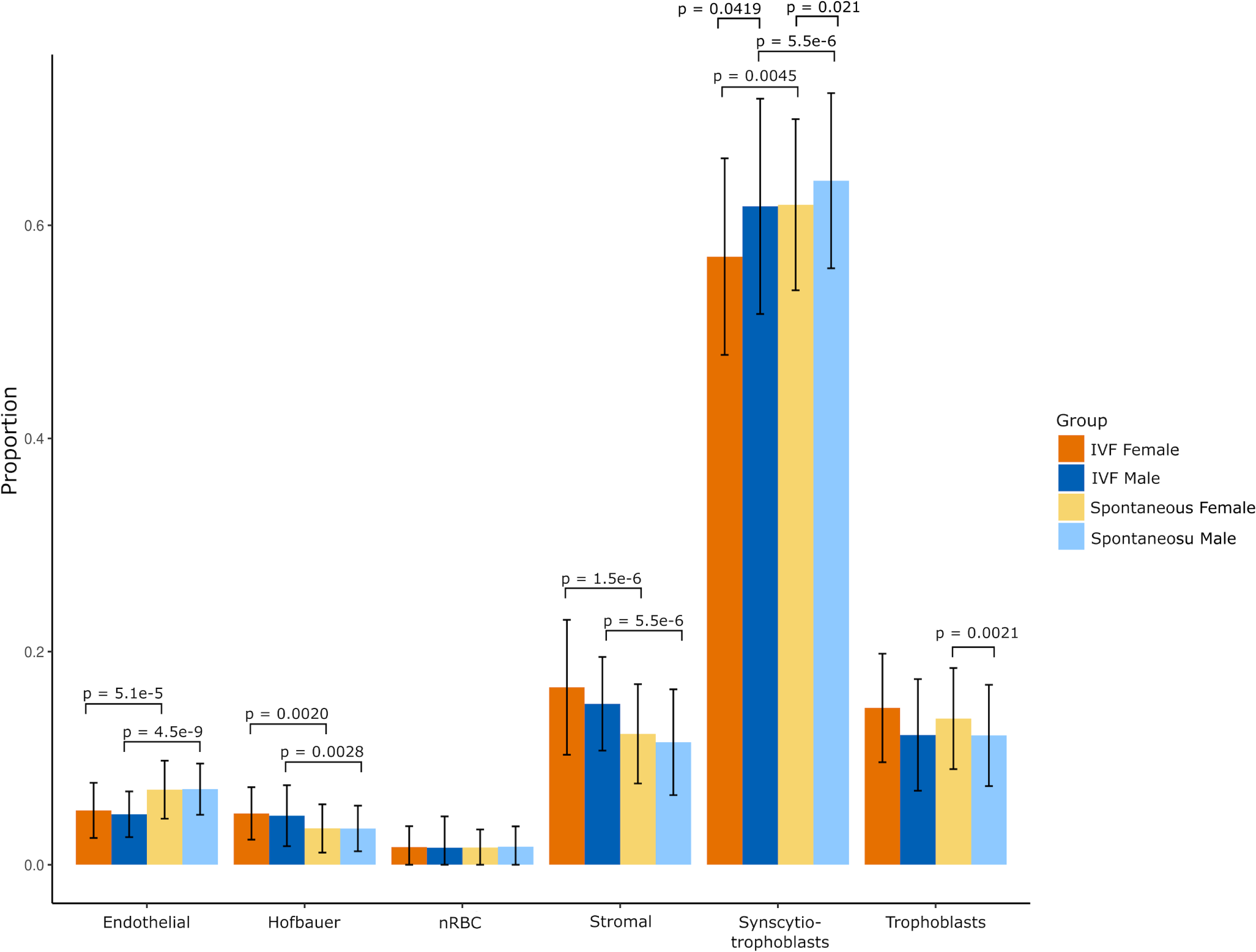
Bar plot depicting mean cell type proportions and standard deviations for each placental cell deconvolution cell type. Significantly different means, calculated using an ANOVA with a Bonferroni test, are indicated above each plot

## Discussion

Incidence of ART has more than doubled over the last 10 years, with IVF as the most common method^7^. Despite its benefits in aiding conception, IVF has been associated with increased incidence of adverse health outcomes such as major birth defects, preeclamapsia, preterm birth and increased cardiovascular risks^13–15^. The IVF protocol typically occurs within the first five days following fertilization^6^ which overlaps with the dynamic epigenetic reprogramming necessary in early preimplantation blastocysts^22^. Therefore, it has been proposed that the IVF protocol and associated stressors may impact early DNA methylation, predisposing the pregnancy to a higher risk of negative health outcomes. The purpose of this study was to investigate the impact of IVF on the placental methylome of healthy, full-term singleton pregnancies using publicly available DNA methylation datasets. We hypothesized there would be differences in DNA methylation in IVF placentae compared to spontaneous controls.

In previous investigations of DNA methylation differences between ART and spontaneous pregnancies, a combination of candidate approaches through pyrosequncing or methylation-specific PCR techniques, and whole genome arrays such as Illumina have been utilized^46–49^. Recent systematic reviews^22,24^ have qualitatively compared the results of studies that analyzed DNA methylation data from ART pregnancies in various tissues, however this remains a complex challenge due to diversity in normalization and statistical approaches across many of the compared studies. Our meta-analysis re-analyzed previously published datasets within a consistent statistical workflow allowing for a more direct and quantitative comparison of sample data.

Screening of datasets on NCBI GEO yielded three studies with raw accessible data that met our inclusion criteria for the meta-analysis. We identified no statistically (FDR <0.05) or differentially methylated (FDR <0.05 and |Δ*β*|>0.05) CpGs between IVF and spontaneous placentae in the mixed fetal sex population. In the female fetal sex stratified population there were no statistically or differentially methylated CpGs between IVF and spontaneous placentae. However, in the male stratified population we identified 9 CpGs that reached statistical significance between IVF and spontaneous placentae. We speculate that our increased ability to observe DNA methylation differences between IVF and spontaneous placentae in sex stratified populations is due to base-line sex-differences in placental DNA methylation that creates a greater spread of DNA methylation intensities when analyzed together, thus reducing statistical significance. In a recent meta-analysis, Andrews et.al. (2022) identified placenta specific differences in DNA methylation due to fetal sex at both site specific and regional levels, further contributing to the argument that analyzing male and female placentae together as a single population may be inappropriate.

Of the 9 statistically significantly different CpGs in IVF males, 3 CpGs were biologically differentially methylated between IVF and spontaneous male placentae. 2 hypermethylated CpGs mapped to the closest transcriptional start sites of *EEF1A2* and *LIPJ*, while 1 hypomethylated CpG mapped to the closest transcriptional start site of *FBRLS1*. *FBRSL1*, a gene involved in RNA binding^51^, was previously found to be differentially methylated in cord blood and blood spots of ART pregnancies^22^. *LIPJ* has previously been found to be significantly enriched in gene sets for gestational diabetes^52^ and has been previously differentially methylated^53^ and differentially expressed^54^ in preeclampsia placentae. *EEF1A2* has shown novel function in angiogenesis through positive feedback regulation via *HIF1A*^55^, an important process in early gestation that is often dysregulated during placental pathologies such as preeclampsia^56^. This is of interest as both gestational diabetes and preeclampsia have been observed at a higher incidence in IVF pregnancies^15,57^. Despite DNA methylation differences at CpGs near these gene regions, further validation in RNA expression studies must be completed to evaluate if the observed methylation differences are contributing to biologically meaningful impacts on transcription in IVF placentae. Beyond mapping to closest transcriptional start sites, it is also important to note that the differentially methylated CpGs closest to *FBRSL1* and *LIPJ* are located between an adjacent enhancer and the gene’s transcriptional start sites. Further gene expression analysis may also identify if differential DNA methylation in these regions impact the enhancer, which may affect nearby genes such as *KCNQ2* and *RNLS* which have both been previously investigated in the context of preeclampsia^58–60^. We also identified 1 statisitcally different CpG on the X chromosome of male placentae in IVF compared to spontaneous that mapped to the closest transcriptional start site of *CAPN6*. Although this site did not reach our threshold to be considered biologically significant, investigation into this gene in the future may be interesting as it is highly expressed in placental tissue and may have various roles in pathology and disease^61^.

Beyond a difference in DNA methylation levels in male IVF placentae compared to spontaneous, we also observed a significant shift to higher levels of DNA methylation variability in both male and female IVF placentae. This agrees with other studies that have also noted higher variability in placental methylation in IVF groups compared to spontaneously conceived populations^49,62^. We further examined the trends of variability in the 3 biologically differentially methylated autosomal CpGs identified in male IVF placentae. While these CpGs did not meet our statistical significance thresholds in females, we saw similar patterns in Δ*β*s and ΔCoVs in females IVF pregnancies compared to the male IVF pregnancies. These differences in DNA methylation variability may partly be explained by the effect of both IVF and sex on relative placental cell populations. Previous reports^50,63^ have reported that differences in placental cell proportions may account for differences and heterogeneity in placental DNA methylation and gene expression. Therefore variability differences observed between DNA methylation in IVF and spontaneous may be driven by differences in cell type composition of the placenta.

The observed trends of DNA methylation and cell composition differences in placental tissue between males and females further highlight how fetal sex may impact placental development and have clinical implications. The placenta matches in sex and genetic characteristics with the developing fetus^64^. Different DNA methylation patterns are observed between male and female placentae which may contribute to placental sexual dimorphism^65^. Therefore considerations of sex may be important to further understand if IVF impacts placental methylation patterns. Hypotheses have previously reported males may adopt strategies of increased investment towards fetal growth and higher placental efficiency, while females may be more responsive to environmental stressors^66,67^. Although there are evident sex differences in placental transcriptomics, there is no indications that support a clear increase in responsiveness to the maternal environment by female fetuses or a dangerous approach utilized by male fetuses^65^. Furthermore, measures of placental efficiency are widely debated, and calculations of efficiency using birth weight:body weight ratios, as reported by Eriksson et.al. (2010), may result in an oversimplification and invalid representation of the true trends of placental function^68^. Further understanding of the implications of this present studies results of differential methylation in male placentae and higher variability in female placentae require validation in animal or cell models to elucidate if these trends are leading to biologically significant differences in placental health or function in a sex specific manor following IVF.

A large hinderance in this study is the lack of robust publicly available data published through open science frameworks such as NCBI GEO with accessibility for re-analysis. It is critical that raw data files with correct formatting are updated frequently to allow for quantitative meta-analyses to validate findings across studies, despite differences in inter-study downstream processing methods. We reccomend those wanting to share biological data, to follow the guidelines as set out by Wilson *et al.* [69]. Further limitations of our study also include a lack of appropriate controls to further understand if these results are likely a result of the IVF protocol or underlying factors of infertility. Additionally, important clinical data such as type of embryo transfer (fresh or frozen) and use of PGT or ICSI were not specified. This highlights that future clinical work in the field of ART should aim to collect more robust data regarding type of ART used, presence of preimplantation genetic testing, frozen or fresh embryo transfer, and fertility status.

To address confounding variables such as fertility status and type of ART used, future work with a mouse study would be beneficial to elucidate if changes in methylation are correlated with IVF protocol involvement or the presence of underlying issues in fertility. Animal models also allow for methylation analysis across gestation, as early methylation differences may contribute to pathology. Additionally, use of animal or cell models may be beneficial to validate if these CpG specific changes in methylation or observed variability may equate to biologically significant impacts in placental development.

In conclusion, the results of this study are critical to further understand the impact of IVF on tissue epigenetics which may help to investigate the connections between IVF and negative pregnancy outcomes. Our study shows that sex specific differences in placental DNA methylation and cell composition should be considered when analyzing placental data, and investigation of mixed-sex population may reduce the computational ability to detect and interpret statistical findings.

## Data availability

All source code can be found on the Wilson Pregnancy Lab Github. All data is already on GEO and how to extract those datasets is within the source code.

## Supporting information

Volcano plot depicting Y chromosome probe methylation differences between IVF and spontaneous placentae in males following linear modelling.

