## Supplementary figures and images for "Examining Fetal Sex-Specific Placental DNA Methylation Intensities and Variability Post *In Vitro* Fertilization"

### Supplemental Data 1

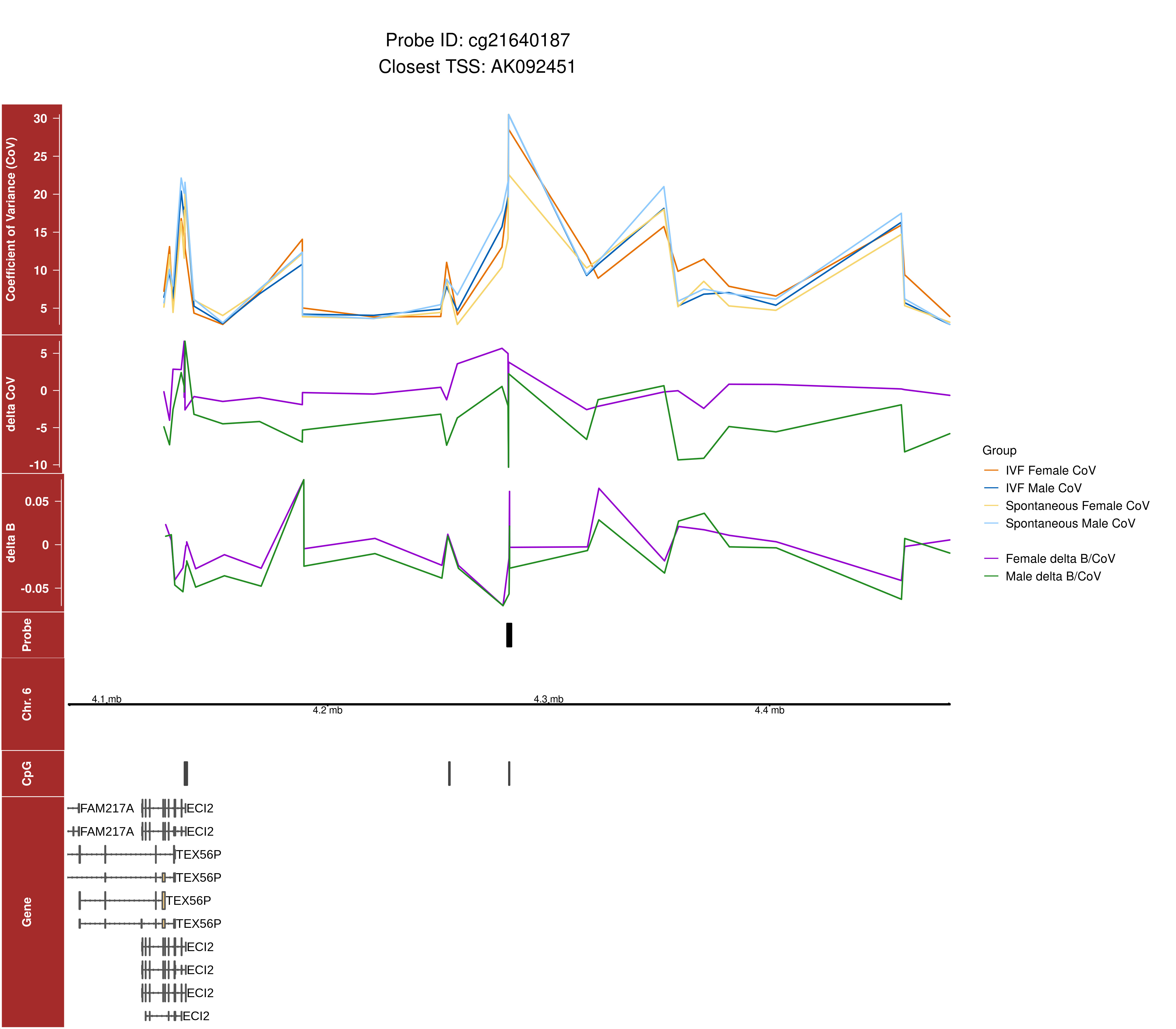

### Supplemental Data 2

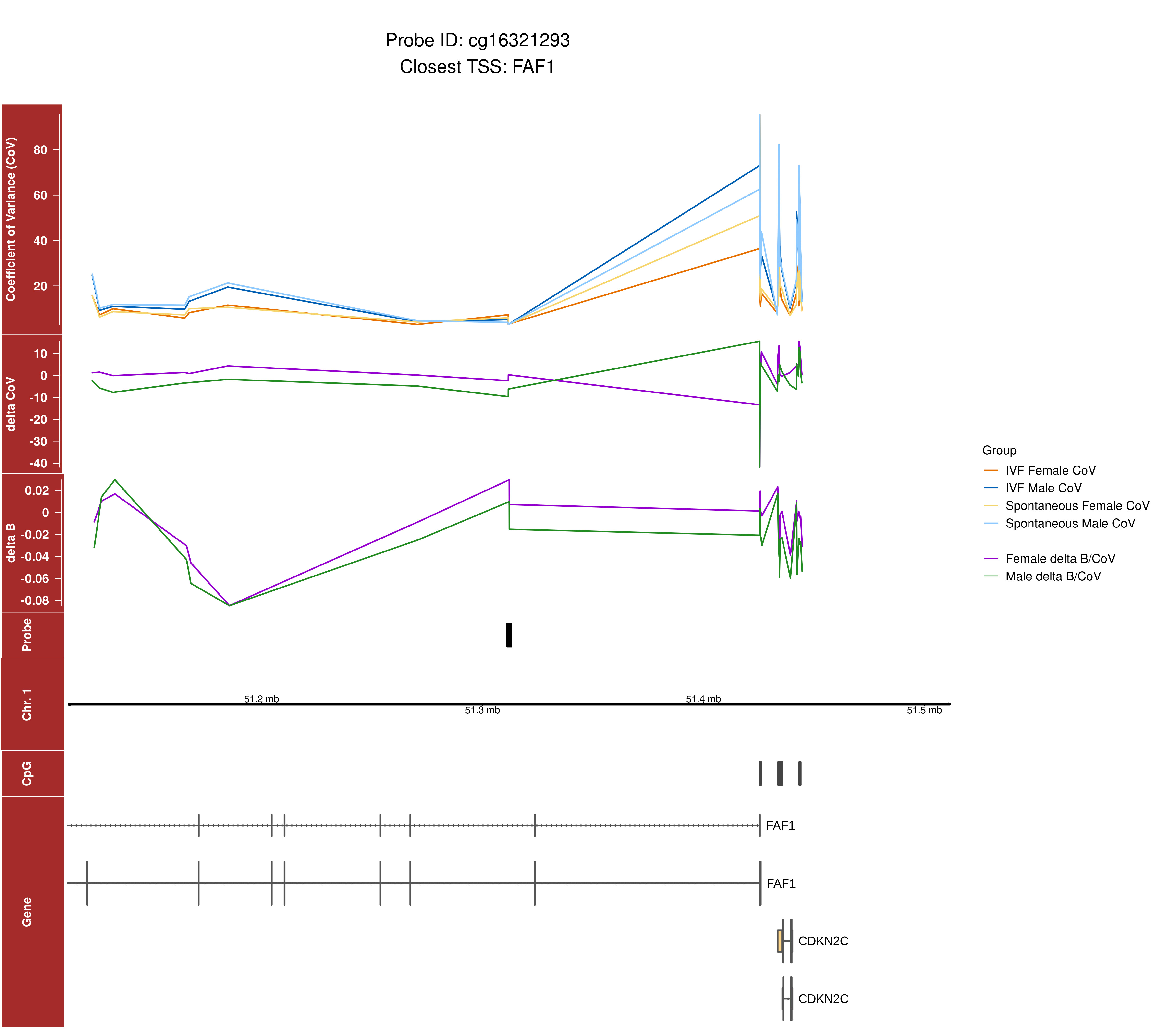

### Supplemental Data 3

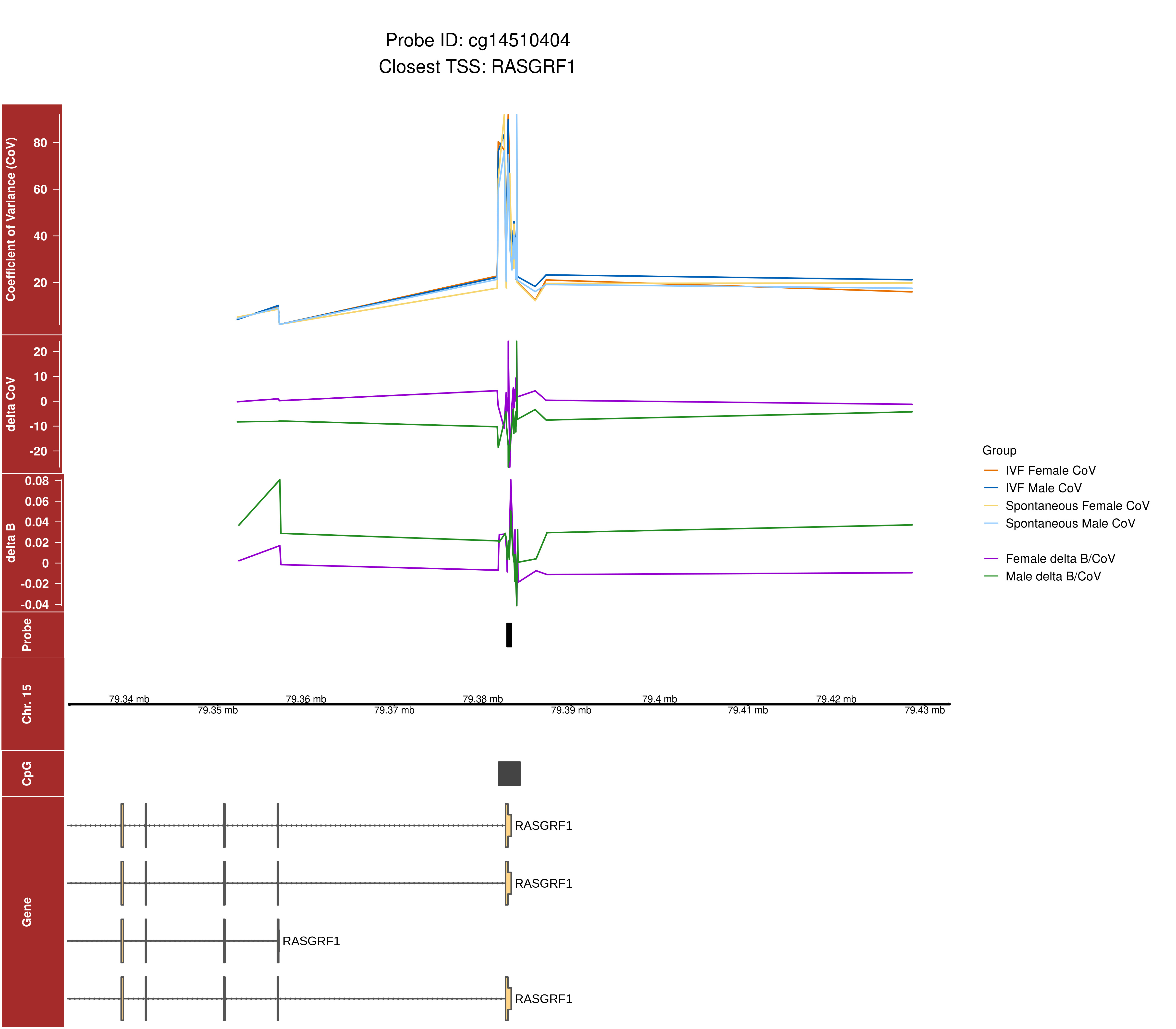

### Supplemental Data 4

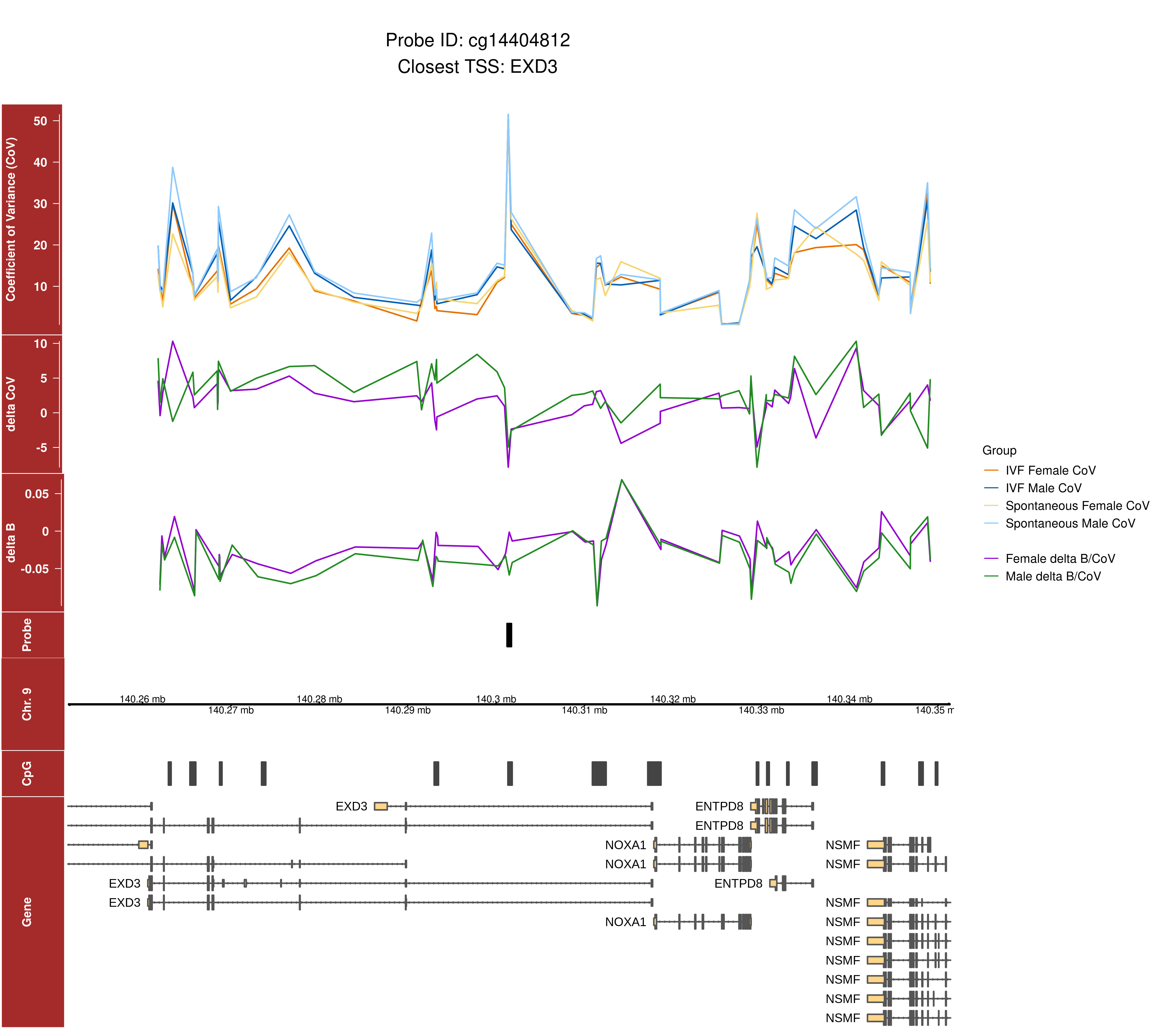

### Supplemental Data 5

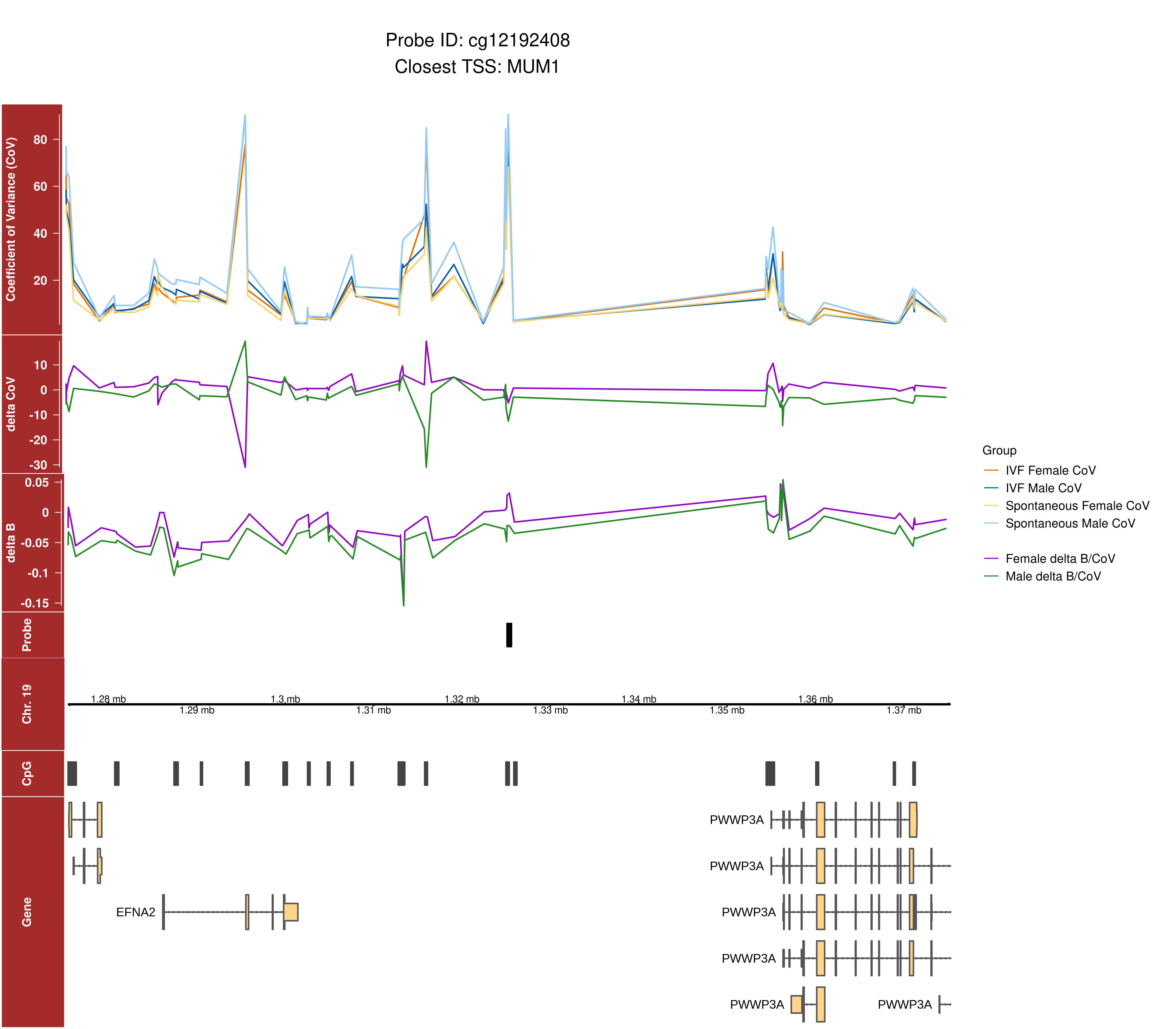

### Supplemental Data 6

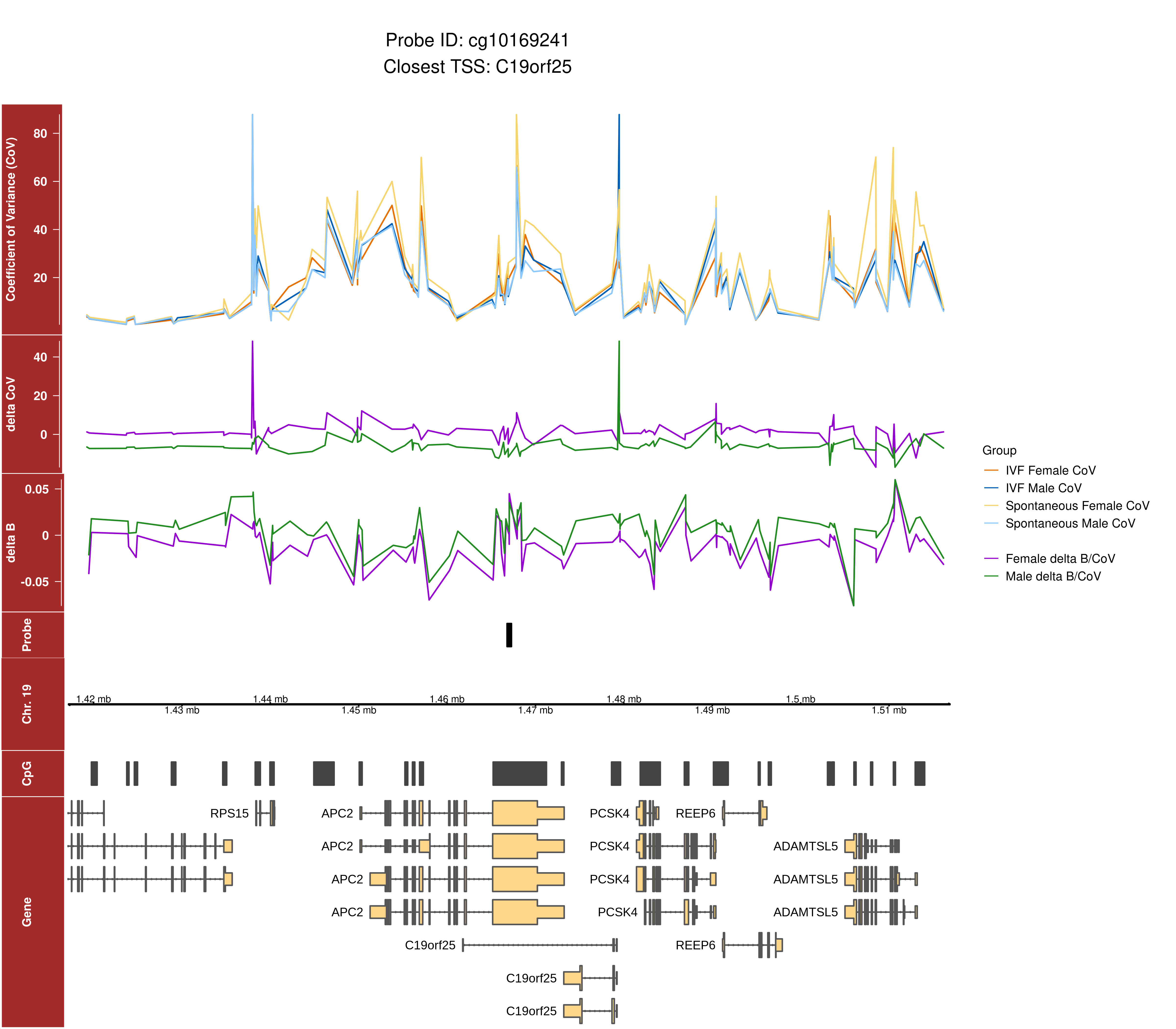

### Volcano plot depicting Y chromosome probe methylation differences between IVF and spontaneous placentae in males following linear modelling.

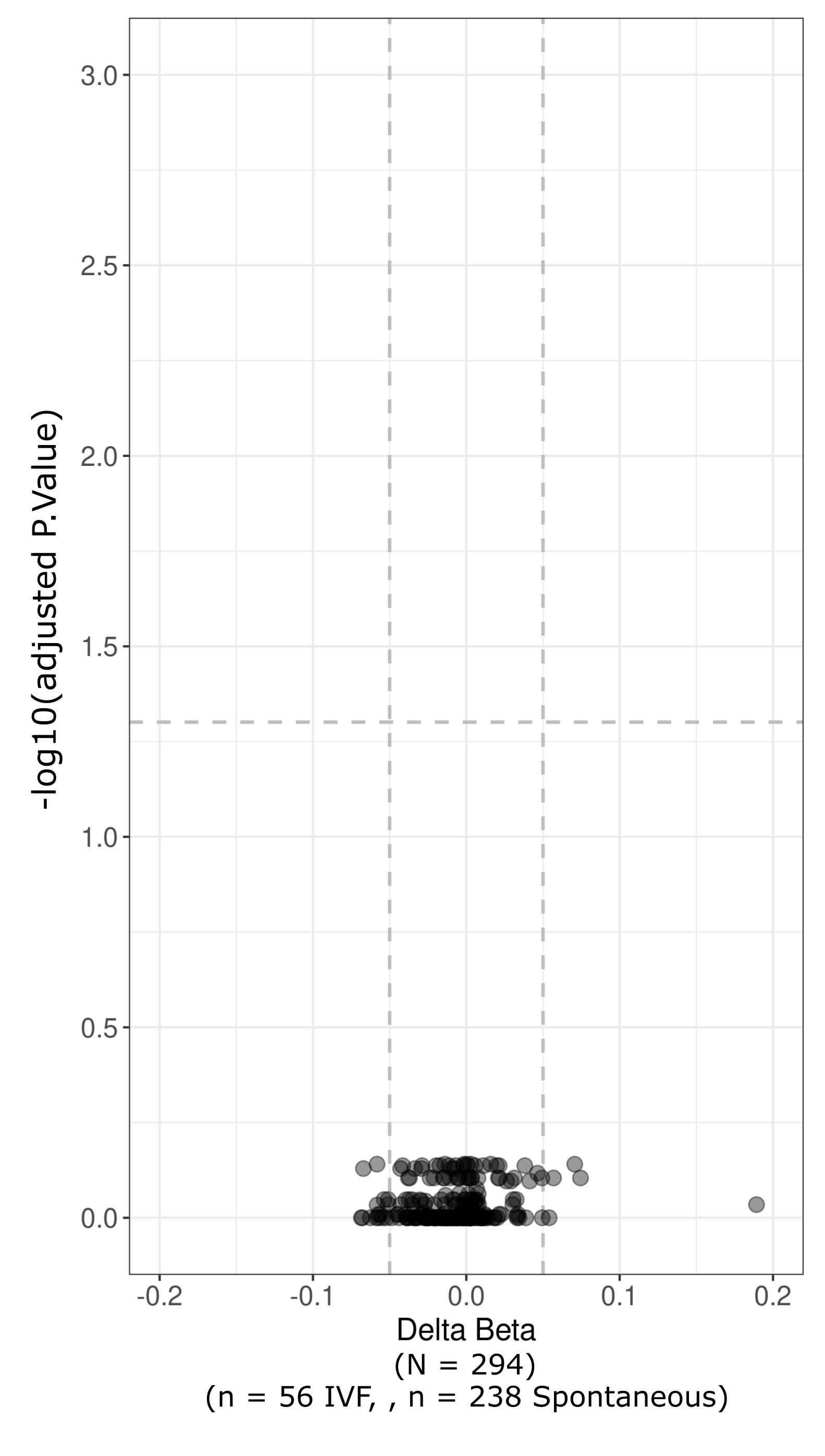
